# Timecourse and convergence of abstract and concrete knowledge in the anterior temporal lobe

**DOI:** 10.1101/2020.06.04.134163

**Authors:** L. Vignali, Y. Xu, J. Turini, O. Collignon, D. Crepaldi, R. Bottini

## Abstract

How is conceptual knowledge organized and retrieved by the brain? Recent evidence points to the anterior temporal lobe (ATL) as a crucial semantic hub integrating both abstract and concrete conceptual features according to a dorsal-to-medial gradient. It is however unclear when this conceptual gradient emerges and how semantic information reaches the ATL during conceptual retrieval. Here we used a multiple regression approach to magnetoencephalography signals of spoken words, combined with dimensionality reduction in concrete and abstract semantic feature spaces. Results showed that the dorsal-to-medial abstract-to-concrete ATL gradient emerges only in late stages of word processing: Abstract and concrete semantic information are initially encoded in posterior temporal regions and travel along separate cortical pathways eventually converging in the ATL. The present finding sheds light on the neural dynamics of conceptual processing that shape the organization of knowledge in the anterior temporal lobe.

## Introduction

How is conceptual knowledge organized, stored and retrieved in the human brain? A “distributed-plus-hub” view of semantic knowledge (Lambon-Ralph, Jefferies, Patterson, & Rogers, 2017; Patterson, Nestor, & Rogers, 2007) suggests that concepts are retrieved through the activation of a network of highly specialized cortical regions that encode information in given sensory modalities (e.g., visual, auditory) or experiential domains (e.g., movement, emotions, human body). During conceptual learning, modality- and domain-specific information converges into a cross-modal hub, identified in the anterior temporal lobe (ATL), where integrated and specific semantic representations are formed. Evidence for this model comes from neuropsychological cases showing that damages in peripheral “spokes” regions often lead to specific semantic impairments in a given modality (e.g., visual, manipulation) or conceptual domain (e.g., tools, colors; Buxbaum, Kyle, Grossman, & Coslett, 2007; Pobric, Jefferies, & Lambon Ralph, 2010; Stasenko, Garcea, Dombovy, & Mahon, 2014); whereas bilateral damages of the ATL lead to a general semantic deficit across modalities and domains of knowledge (Guo et al., 2013; Hodges, Patterson, & Tyler, 1994; Rogers, Ralph, Hodges, & Patterson, 2004).

A recent update of the distributed-plus-hub model (Lambon-Ralph et al., 2017) put forward the idea that semantic representations in the ATL hub are organized according to a dorsal-to-medial and abstract-to-concrete gradient: Whereas the representation of concrete features insists on the medial-ventral ATL, abstract features are represented in the dorsal-lateral ATL. Empirical support for a graded ATL hub comes from functional magnetic resonance imaging (fMRI) studies comparing abstract and concrete concepts (Hoffman, Binney, & Lambon Ralph, 2015; Striem-Amit, Wang, Bi, & Caramazza, 2018). However, other fMRI studies following a similar methodology (including one meta-analysis) failed to find evidence for such an organization of semantic knowledge in ATL (Binder, Westbury, McKiernan, Possing, & Medler, 2005; Wang, Conder, Blitzer, & Shinkareva, 2010). One reason for this discrepancy may lie in the fact that the ATL is shy to fMRI, due to the drop of BOLD signal near air cavities, calling for confirmatory results using alternative methodologies. Moreover, several questions about the role of ATL in conceptual processing remain unanswered. For instance, it is unclear at which stage of conceptual retrieval semantic representations emerge as a gradient in the ATL. Indeed, previous chronometric tests focused alternatively on concrete (Borghesani & Piazza, 2017; Chan et al., 2011; García et al., 2019; Jackson, Lamobon Ralph, & Probic, 2015; Mollo et al., 2017; Teige et al., 2019) or abstract aspects of word meaning (Fahimi Hnazaee, Khachatryan, & Van Hulle, 2018), and failed to show a specific ventral or dorsal ATL activity related to concrete and abstract features.

A related question is why semantic information is organized in the ATL according to a dorsal-to-medial and abstract-to-concrete gradient. One possibility is that this pattern depends on the long-range connectivity profile of different subparts of the ATL (Lambon-Ralph et al., 2017). According to this hypothesis, the medial-ventral ATL responds more to concrete concepts by virtue of having greater connectivity to visual areas through the ventral occipital-temporal cortex (VOTC); whereas the dorsal-lateral ATL contributes more to abstract concepts by virtue of its greater connectivity with the posterior temporal language system and with orbito-frontal regions that support social cognition and emotional value (see Figure 3A for a depiction of this connectivity model by Lambon-Ralph and collaborators). This model is partially supported by tractographic studies (Binney, Parker, & Lambon Ralph, 2012; Chen, Lambon Ralph, & Rogers, 2017) and functional connectivity analysis during rest (Jackson, Hoffman, Pobric, & Lambon Ralph, 2016; Pascual et al., 2015) showing the presence of this cortico-cortical tracks in human subjects. However, there is no direct evidence that concrete and abstract information travels between peripherical spokes and subparts of the ATL-hub, along these cortical routes, during the retrieval of specific concepts.

**Figure 1.**
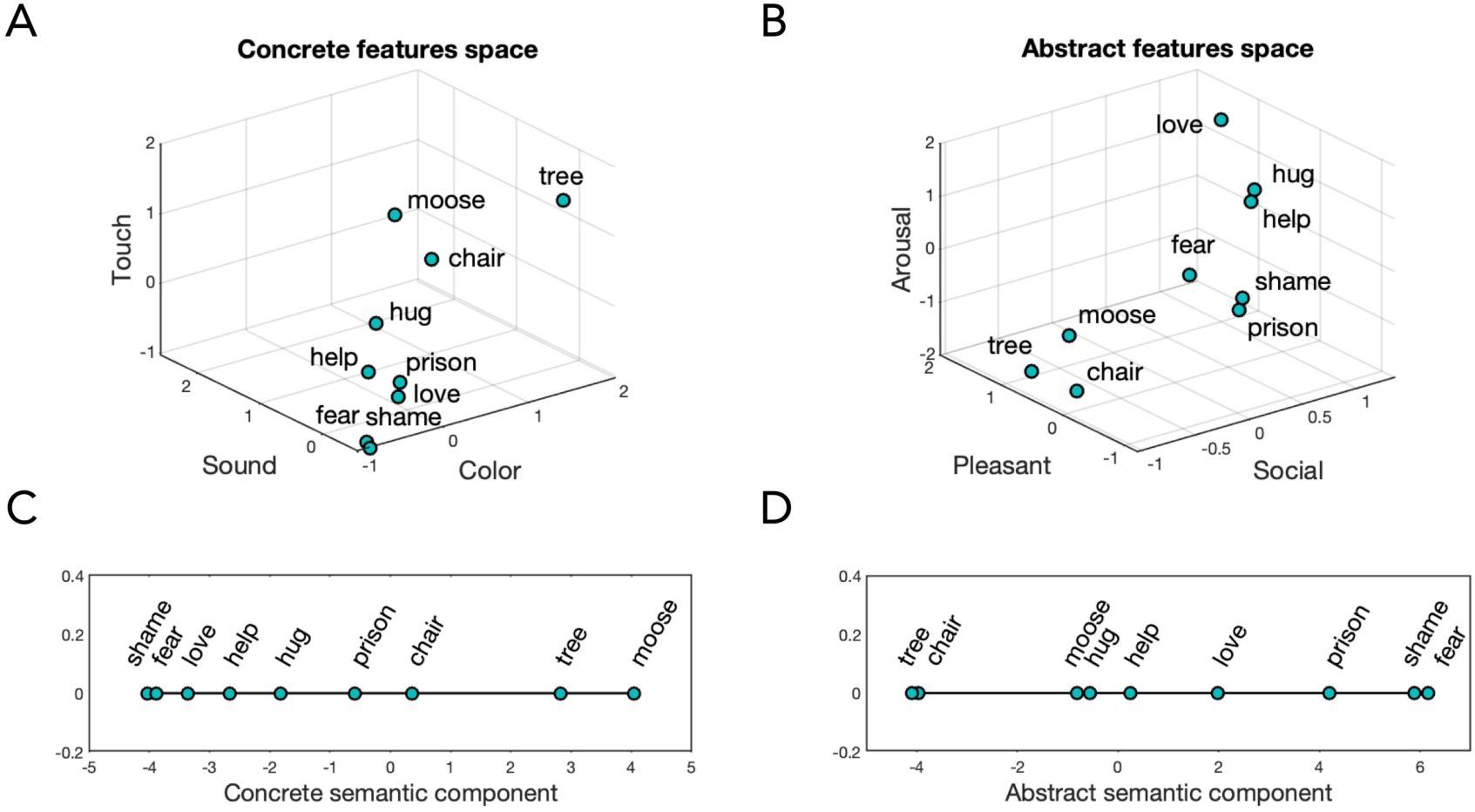
Dimensionality reduction. A) Schematic representation of a 3-D semantic space where each word is viewed in a coordinate system defined by concrete features such as Touch, Sound and Color (the actual multidimensional space comprised 31 dimensions, here reduced to 3 for visualization purposes). B) Schematic representation of a 3-D semantic space where each word is viewed in a coordinate system defined by abstract features such as Arousal, Pleasant and Social (the actual multidimensional space comprised 31 dimensions). C) Words’ weights along the first principal component of the concrete space. D) Words’ weights along the first principal component of the abstract space.

**Figure 2.**
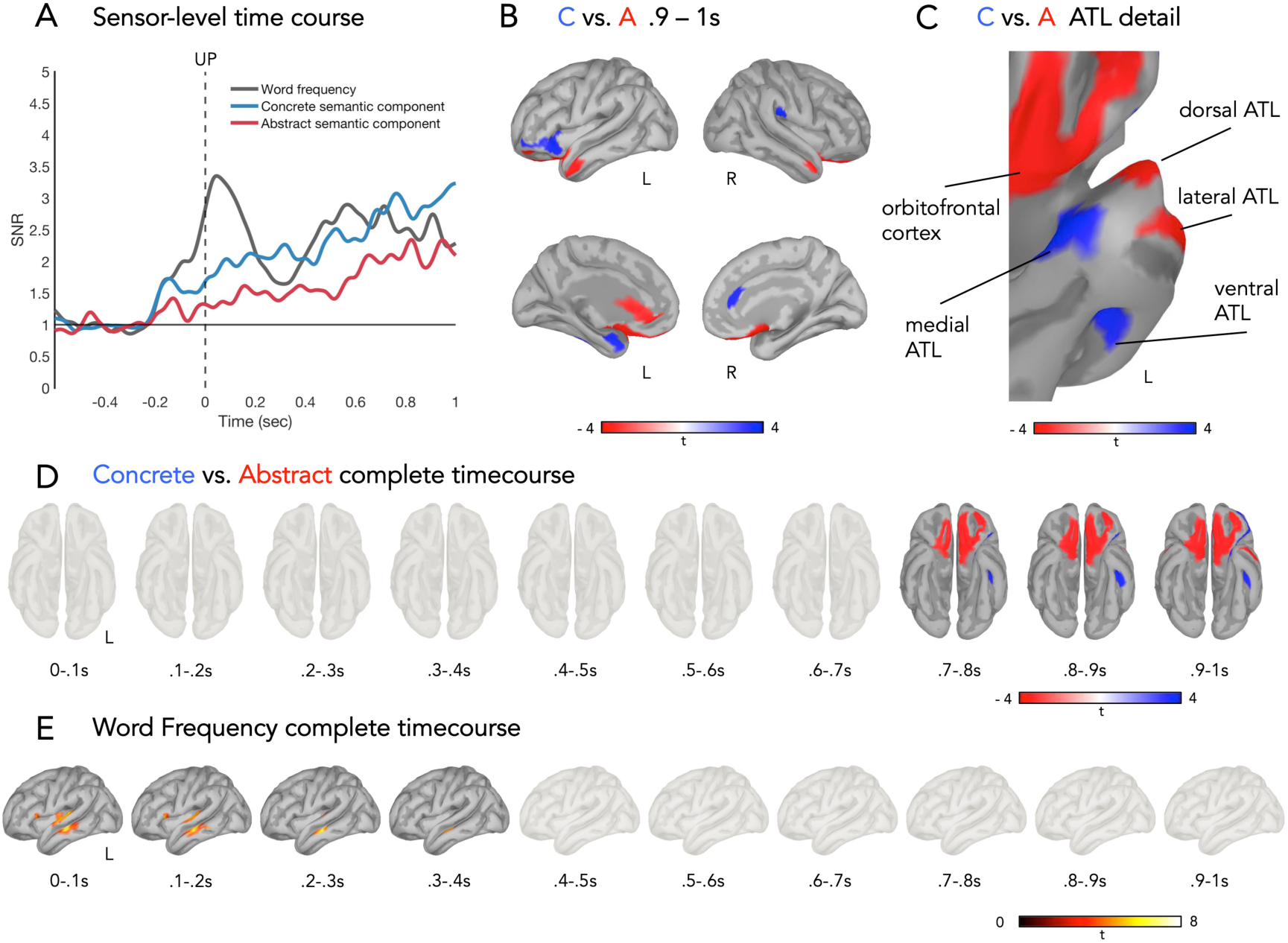
Spatiotemporal dynamics of lexical and conceptual representations. A) Root-mean-square of the SNR of ERRC of the word frequency (grey), concrete semantic component (blue) and abstract semantic component (red) predictors. 0s = uniqueness point. B) Concrete > Abstract (one-sample t-test (two-tailed), FDR-corrected *p* < .05, > 15-vertex) in the .9 to 1s interval. C) Detail on the left ATL for the contrast Concrete > Abstract (one-sample t-test (two-tailed), FDR-corrected *p* < .05, > 15-vertex) in the .9 to 1s interval. D) Source-reconstructed statistical maps of the contrast Concrete > Abstract (one-sample t-test (two-tailed), FDR-corrected *p* < .05, > 15-vertex) in consecutive 100ms intervals. E) Source-reconstructed statistical maps of the Word Frequency predictor (one-sample t-test (one tail), FDR-corrected *p* < .05, > 15-vertex) in consecutive 100ms intervals.

**Figure 3.**
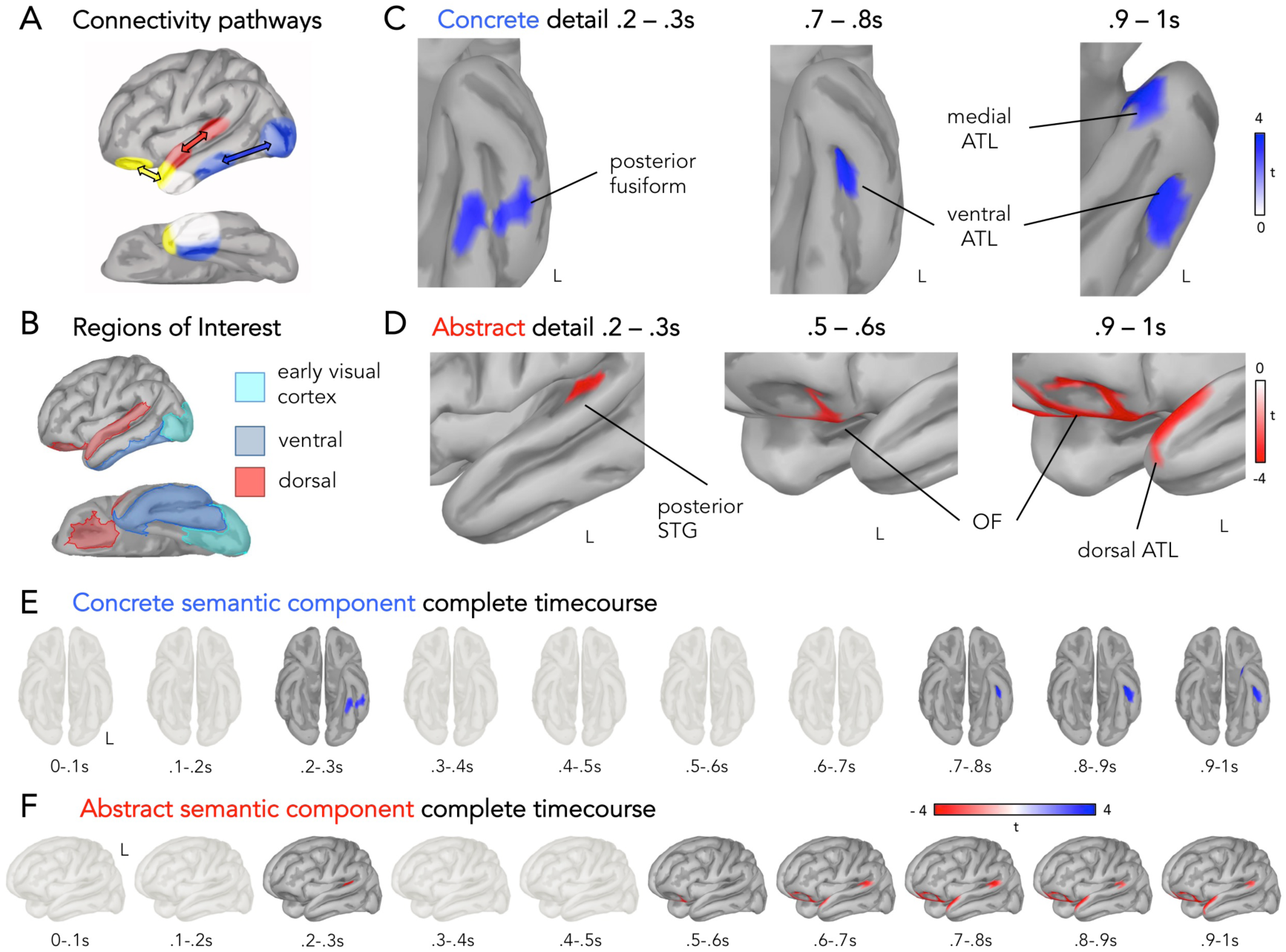
A dorsal and a ventral stream. A) Neuroanatomical sketch of hub and spokes connectivity pathways (see also Lambon-Ralph et al., 2017). B) Dorsal (red) ventral (blue) and early visual cortex (cyan) ROIs in the left hemisphere. C) Concrete semantic component activation detail on the left ventral and medial temporal cortex in the relevant time intervals (one-sample t-test (two-tailed), FDR-corrected *p* < .05, > 15-vertex). D) Abstract semantic component activation detail on the left posterior STG and left ATL in the relevant time intervals (one-sample t-test (two-tailed), FDR-corrected *p* < .05, > 15-vertex). E) Source-reconstructed statistical maps of the contrast Concrete > Abstract (one-sample t-test (two tailed), FDR-corrected *p* < .05, > 15-vertex) in consecutive 100ms intervals, positive t-values. F) Source-reconstructed statistical maps of the contrast Concrete > Abstract (one-sample t-test (two-tailed), FDR-corrected *p* < .05, > 15-vertex) in consecutive 100ms intervals, negative t-values.

In the present magnetoencephalography (MEG) study we aim to investigate the spatiotemporal organization of semantic knowledge in the brain. In particular, we will focus on abstract and concrete semantic information encoding in the attempt to: (i) assess whether and when a semantic gradient emerges in the ATL and (ii) shed some light on how the information concerning abstract and concrete conceptual dimensions reaches anterior temporal brain regions. To this end we recorded MEG signals from thirty participants performing a semantic categorization task on 438 spoken words. Each word referred to a concept (e.g., chair, dog, policeman) that was independently rated across 65 feature dimensions (e.g., color, shape, happiness, arousal, cognition, etc.; Binder et al. 2016). Thus, each word could be considered as a point in a high-dimensional feature space. Principal component analysis (PCA) was implemented in order to reduce dimensionality and create high-level abstract and concrete semantic predictors. We then used a combination of multiple linear regressions analysis and source reconstruction methods to assess the spatiotemporal dynamics of abstract and concrete semantic information processing.

## Results

### Behavioral results

Participants listened to auditory-presented words and were instructed to categorize each stimulus as either related to sensory perception (i.e., they refer to something that can be easily perceived with the senses, like “red” and “telephone”), or unrelated to sensory perception (i.e., they refer to something that cannot easily be perceived with the senses, like “agreement” and “shame”). We expected participants to categorize relatively concrete words as related to sensory perception and relatively abstract words as unrelated to sensory perception. To assess this, we correlated participants’ responses with concreteness estimates for each item (i.e., the concrete principal component, see below). The results indicated a significant association between participants’ responses and concreteness estimates (r(436)=.82, *p* < .001). We did not analyze reaction times because participants’ responses were delayed in order to avoid motion-related artifacts in the MEG signal (i.e., see Material and Methods for details).

### Mapping sound to meaning

Spatiotemporal dynamics of conceptual retrieval were inferred using multiple linear regression analysis of MEG data (Chen, Davis, Pulvermüller, & Hauk, 2013; Hauk, Davis, Ford, Pulvermüller, & Marslen-Wilson, 2006; Hauk, Pulvermüller, Ford, Marslen-Wilson, & Davis, 2009; Miozzo, Pulvermüller, & Hauk, 2015). We focused on three predictors spanning both lexical and semantic aspects of word retrieval (word frequency, and the abstract and concrete semantic predictors that were computed via PCA, see below). Also, we included in the model other predictors to control for potentially confounding variables (i.e., word duration and participants’ judgment of each item as related [1] or unrelated [0] to the senses).

Word frequency was calculated as the frequency of occurrence of a given word in a large corpus of text samples (subtlex-it, Crepaldi et al., 2013). Semantic predictors were derived instead from Binder’s et al., (2016) database. As briefly mentioned above, each word in Binder’s work was rated across 65 fundamental semantic features. Some of these features were related to sensory experience (e.g., sound, shape, smell), whereas others to social, emotional or intellectual experiences (e.g., arousal, social, sad). Following the concrete versus abstract labeling provided in the original database (Binder et al., 2016), we separated the entire semantic space (65-Dimension) into concrete (31-Dimensions) and an abstract (31-Dimensions) features (with three feature dimensions discarded for missing values; See Methods). Thus, each word could be considered as a point in a concrete semantic space (see Figure 1A), and in an abstract semantic space (see Figure 1B). We used principal component analysis (PCA) to reduce the dimensionality of the dataset and adopted the first concrete (Figure 1C) and the first abstract semantic component (Figure 1D), to represent the same data in a new one-dimensional coordinate system. Importantly, the resulting semantic components do not simply reflect how concrete and how abstract a word is, but instead represents concrete and abstract aspects of concepts in a new low-dimensional space that encodes the most salient structural features of the high-dimensional space from which it is derived. For instance, in the concrete principal component, “moose” is more similar to “chair” than to “hug”, whereas the opposite is true in the abstract principal component.

Access to any word’s lexical-semantic properties obviously depends on the unique identification of that word (Marlsen-Wilson, 1987). Therefore we aligned our multiple regression analysis to the uniqueness point of each word (UP), that is, the point in time when the acoustic and phonetic information already presented (e.g., the syllables “ba”-”nan”) is compatible with a single lexical entry (i.e., banana). Thus, for each time point, channel and subject we calculated event-related regression coefficients (ERRCs) reflecting the contribution of each predictor to the MEG signal. The spatiotemporal dynamics of the different predictors were characterized as the root-mean-square (RMS) of the signal-to-noise ratio (SNR) of ERRC (see Material and Methods). This provided a unified measures of sensor-level activity (magnetometers and gradiometers are combined together; Figure 2A). Source-reconstructed statistical maps of single predictors were computed as consecutive 100ms average time-windows (Figure 2B/C/D/E). The same analysis, time-locked to the word onset, is reported in the supplementary materials and confirms the results described here (Supplementary Figure 1).

Consistent with an optimally efficient recognition system, lexical access occurred shortly after the UP, essentially validating our UP estimation procedure (see also Kocagoncu, Clarke, Devereux, & Tyler, 2016). This is depicted in Figure 2A by the peak in the SNR of the word frequency predictor, from 0 to 300 ms (with zero being the uniqueness point of each word). Dual-stream models of speech processing state that, at this point in time, the spectrotemporal and phonological analysis of speech signals maps onto lexical representations stored in middle temporal regions (Hickok & Poeppel, 2007). Accordingly, in our study, encoding of word frequency information involved bilateral middle temporal regions, the superior temporal gyrus (STG) and the inferior frontal gyrus (IFG), with an overall weak left-hemisphere bias (Figure 2E). At later time windows, the word frequency predictor was encoded in right premotor areas, right medial temporal lobe, left anterior temporal lobe and left parahippocampal formation (Supplementary Figure 2).

Semantic information encoding occurred at later time stages. Specifically, abstract and concrete semantic components showed a sustained increase in the SNR starting from 300ms after the UP and continuing until the end of the trial (Figure 2A). Consistently, a recent electroencephalography (EEG) study reported effects of concreteness, during spoken word recognition, in a similar time window (i.e., 400 - 900ms; Winsler, Midgley, Grainger, & Holcomb, 2018).

One important prediction of the distributed-plus-hub model (Lambon-Ralph et al., 2017) is that semantic representations in the ATL follow a dorsal-to-medial and abstract-to-concrete gradient. To test this, we contrasted source-reconstructed ERRCs of the concrete and abstract semantic components in consecutive 100ms time windows. Figure 2D shows that, at late latencies (i.e., > 700ms), anterior temporal regions responded to both types of semantic information. Crucially, semantic information encoding followed a dorsal-to-medial, abstract-to-concrete gradient. This is illustrated in greater detail in Figure 2C where in the 900ms to 1000ms time window, concrete semantic encoding engaged the left ventral and medial ATL and abstract semantic encoding involved the anterior lateral ATL and the left dorsal ATL. Additionally, the left IFG, right supramarginal gyrus (SMG) and right medial prefrontal cortex (MPFC) responded preferentially to concrete semantic information (Figure 2B). Conversely, bilateral orbitofrontal cortex (OF) and left MPFC responded preferentially to abstract semantic information (Figure 2B). Different parts of the ATL (i.e., dorso-lateral and ventral-medial), therefore, seem to integrate within two different networks of regions implicated in the representation of abstract and concrete features, respectively.

### Streams of semantic information

We hypothesized that semantic information travels along parallel paths (one dorsal-abstract and one ventral-concrete) to reach the ATL. To test this hypothesis, we increased the sensitivity of our analysis (by reducing the number of multiple comparisons for which correction is required) and adopted a region of interest (ROI) approach. That is, statistical analysis of abstract and concrete semantic components was restricted to three macro-regions: Early visual cortex (EVC), ventral occipito-temporal cortex (VOTC; henceforth, ventral ROI) and a macro region including the superior temporal gyrus and the orbito-frontal cortex (Figure 3B, see Material and Methods for details). Our choice of ROIs was motivated by current models of long-range ATL connectivity pathways (see Figure 3A; Lambon-Ralph et al., 2017). Moreover, we separated the EVC from the anterior ventral occipital cortex due to the fact that these two macro-regions have different roles during semantic processing (Bracci & Op de Beeck, 2016; Clarke & Tyler, 2014; Mattioni et al., 2020) as well as to balance the number of vertices across different ROIs (see Method sections for details).

The time course of concrete semantic information encoding is illustrated in Figure 3E. The EVC ROIs did not reveal any significant differences between abstract and concrete semantic information encoding. The first observable response was confined to left ventral ROI. An enlarged view of this portion of the temporal lobe in the time window of significance (i.e., 200ms to 300ms, Figure 3C) shows a cluster of activation in the posterior fusiform and mid-lateral temporal regions. Notably, encoding of concrete semantic information progressively engaged more anterior regions. Specifically, at 700ms after word recognition, concrete information was encoded in left ventral ATL and at 900ms in left medial ATL (Figure 3C).

With a similar time course, abstract semantic information encoding progressed from posterior to anterior portions of the left dorsal ROI (Figure 3F). Figure 3D depicts the earliest (200ms to 300ms) observable response to abstract semantic information localized in the left posterior STG. Importantly, the OF cortex responded only at later time intervals (> 500ms; Figure 3D), immediately before the dorsal ATL (> 600ms), and all these areas remained active until the end of the trial (Figure 3D; see also movies A and B).

The overall pattern of results strongly suggests the existence of two streams of semantic processing. Both abstract and concrete semantic information encoding progressed from posterior to anterior temporal regions. That is, along the left ventral stream, concrete semantic information initially involved the fusiform cortex, followed by left ventral ATL and medial ATL responses. Moreover, along the left dorsal stream, abstract semantic information involved, at subsequent time intervals, STG, OF, and anterior dorsal ATL.

## Discussion

To test whether and when an abstract-to-concrete graded semantic representation emerges in the ATL and how does conceptual information reach anterior temporal brain regions we took advantage of the high spatiotemporal resolution of MEG signals. Using a multiple linear regression analysis of MEG-recorded brain activity we obtained for every time point, channel and subject event-related regression coefficients (ERRC) reflecting the contribution of each predictor to the data. Predictors of interest included variables associated with the frequency of each word as well as variables related to abstract and concrete dimensions of semantic knowledge. Sensor-level results showed sequential encoding of lexical and semantic information (see Figure 2A), suggestive of a serial organization of spoken word recognition (for similar findings see Brodbeck, Presacco, & Simon, 2018; Winsler et al., 2018). Furthermore, in line with prominent models of speech processing (Hickok & Poeppel, 2007), source-reconstructed cortical responses to the different predictors evidenced a sound-to-meaning mapping system that includes middle and superior temporal areas bilaterally (Supplementary materials Figure 1), and culminates in concrete and abstract semantic representations being encoded in the ATL. Crucially, left-hemispheric ventral and medial ATL regions responded preferentially to concrete aspects of conceptual knowledge. More abstract features, instead, were encoded in the anterior dorsal and lateral ATL areas.

We then moved on to investigate how abstract and concrete semantic information reaches the ATL during conceptual retrieval. Using a region of interest approach motivated by current models of long-range ATL connectivity pathways (see Figure 3A), we showed that the earliest observable responses to concrete and abstract semantic information laid, respectively, in the fusiform gyrus and in the posterior STG. Moreover, whereas concrete semantic information sequentially involved more anterior areas along the ventral temporal lobe, abstract semantic processing involved, at later time stages, the orbitofrontal cortex and the dorsal anterior temporal lobe.

### The role of the ATL hub in conceptual processing

To date, empirical evidence for a graded ATL hub mainly stems from fMRI studies comparing abstract and concrete concepts (Hoffman et al., 2015; Striem-Amit et al., 2018). Due to the low temporal resolution of fMRI techniques, however, the time course of graded semantic representations in the ATL is still largely unknown. The present MEG study shows that the ATL gradient emerges at late time stages of spoken word recognition, with sustained neuronal response between 600 and 1000ms after lexical access (Figure 2C). This late temporal window suggests that the graded activity of the ATL may be involved in the integration of different features of a retrieved concept into a cohesive construct, a role often attributed to the ATL-hub (Clarke & Tyler, 2015; Coutanche & Thompson-Schill, 2015; Lambon-Ralph et al., 2017; Pylkkänen, 2019). Such a role is also supported by our finding that abstract and concrete features are encoded in different brain regions at earlier time windows and seem to converge into the ATL along separate cortical pathways. Moreover, this timecourse is in line with previous chronometric studies of semantic processing in the ATL using electrocorticography recordings (Chen et al., 2016), TMS (Jackson et al., 2015) and MEG (Borghesani, Buiatti, Eger, & Piazza, 2018), which show that the ATL starts to encode semantic information in a sustained way from ∼300ms after stimulus onset (in the case of visually presented stimuli, which are usually processed faster and less incrementally than auditory spoken words; Vartiainen, Parviainen, & Salmelin, 2009), until 800-1000ms after stimulus onset.

It is important to note, however, that some studies have reported transient semantic effects in the ATL as early as ∼100ms after stimulus onset (Chan et al., 2011; Mollo et al., 2017; Teige et al., 2019), and an opposite flow of semantic information, from the ATL-hub to the spokes, has been suggested (Mollo et al., 2017). However, this early ATL activity has been found only in the case of gross categorical distinction (Borghesani et al., 2018; Chan et al., 2011; Teige et al., 2019) and it is almost invariably followed by sustained ATL activations occurring between ∼300 and 1000ms after stimulus onset (Borghesani et al., 2018; Chan et al., 2011; Mollo et al., 2017; Teige et al., 2019). This state of affairs opens to the intriguing possibility that the ATL may transiently encode superordinate categorical distinctions (Clarke, Devereux, Randall, & Tyler, 2015; Clarke, Taylor, Devereux, Randall, & Tyler, 2013) that facilitate the activation of the relevant spokes (Borghesani et al., 2018; Chiou & Lambon Ralph, 2019; Mollo et al., 2017), which in turn feed-back detailed domain-specific information for a full-fledged semantic access to specific concepts (Clarke & Tyler, 2015; Lambon-Ralph et al., 2017).

However, an early and transient semantic activity in the ATL has been reported only in experiments using images (Clarke et al., 2015, 2013), visually presented words (Borghesani et al., 2018; Chan et al., 2011; Teige et al., 2019) or words and images together (Mollo et al., 2017), and may be specific for the visual modality. In the case of visual stimuli (especially in the case of images) the early activity in the ATL may be the consequence of an automatic feedforward sweep of neural responses through occipital and ventral temporal cortices (Chen et al., 2016; Clarke & Tyler, 2015; Rupp et al., 2017) that would be absent in the case of auditory stimuli, as in the present experiment. Indeed, explorative analysis of our data did not provide convincing evidence for an early ATL activity related to superordinate gross categorical representations (Supplementary Figure 3C), although this result cannot be taken as conclusive.

In sum, although the ATL might be involved in semantic processing at different levels (superordinate gross distinction, specific multimodal concepts) and in different timepoints (early, late activations), our results suggest that the graded organization of abstract and concrete conceptual features in the ATL emerges in the late stages of conceptual processing, as the product of convergent conceptual information from different cortical streams, and possibly coinciding with the retrieval of a cohesive and specific conceptual representation.

### Routes to meaning in the brain

After an initial sound to meaning mapping in the middle temporal gyrus (MTG), signaled by a strong sensitivity to the frequency component right after the word uniqueness point, abstract and concrete semantic information starts to be encoded in two different temporal regions, one above and one below the MTG. Brain activity associated with concrete semantic features emerges sequentially in the fusiform gyrus, the ventral ATL and the medial ATL. Neural correlates of abstract semantic knowledge follow a parallel dorsal path: posterior STG, OFC and finally the dorsal-lateral ATL.

Previous studies have provided evidence for long-distance connections between these brain regions and different subparts of the temporal pole (Binney et al., 2012; Chen et al., 2017). The discovery of these white-matter tracks suggested that the gradient-like organization of the ATL is due to the fact that sensory, emotional and linguistic information travels along different cortical pathways during conceptual learning, and is stored in different ATL regions situated at the termination of these pathways. Once the concept is stored in the ATL, it could in principle be directly reactivated without going through these cortical pathways again (Fairhall & Caramazza, 2013; Mollo et al., 2017). Instead, our chronometric data offer the first direct evidence that these pathways are at least partially retraced during conceptual retrieval. This result suggests that semantic knowledge in the ATL may emerge through the contribution of the entire network, instead of a simple re-activation of stand-alone representations. The re-instantiation of this generative process during conceptual retrieval may be useful to allow a constant representational update in long-term memory, and provide the flexibility to retrieve a given concept highlighting specific feature dimensions instead of others. Once again, this position confirms the central role of the ATL in conceptual retrieval, as suggested by several neuropsychological studies (Abel et al., 2015; Chen et al., 2016b; Coutanche & Thompson-Schill, 2015; Hoffman et al., 2015; Striem-Amit et al., 2018).

The present findings corroborate the existence of two parallel cortical streams for concrete and abstract semantics travelling along dorsal and ventral temporal areas and ultimately terminating in the ATL. This result, however, does not exclude the existence of other routes. For instance, it is conceivable that sensorimotor information, mostly encoded in parietal and motor areas (Fernandino et al., 2015; Pulvermüller, Shtyrov, & Ilmoniemi, 2005), would reach temporal and inferior frontal areas travelling along the arcuate fasciculus (Motivating the IFG modulation by the concrete semantic regressor; Pulvermüller, 2013). Alternatively, evidence from a recent tractography study (Chen et al., 2017) has suggested two possible routes connecting the ATL to the parietal cortex via the posterior middle temporal gyrus and posterior fusiform gyrus. An important open question for future research therefore concerns the relative functional contribution of different streams of information to conceptual representations.

### Extended networks involved in the representation of concrete and abstract features in the brain

The representation of abstract and concrete semantic knowledge extends also beyond the ATL and the dorsal/ventral pathways in the left hemisphere. Abstract information additionally engages the right OFC and the left mPFC (see Sabsevitz, Medler, Seidenberg, & Binder, 2005; Wang et al., 2018; for similar results). On the other hand, the concrete regressor activated a different network of regions that are typically involved in the representation of multimodal concrete knowledge (Binder, Desai, Graves, & Conant, 2009; Binder et al., 2005; Fernandino et al., 2015) and included the left IFG, right SMG, and right mPFC. Moreover, removing the constraint originally imposed to the cluster size (>15 vertices), but keeping a conservative statistical threshold (FDR-corrected *p* < .05 at the whole-brain level), concrete-related activity emerges also in the left angular gyrus, which is considered an important multimodal hub for the integration of concrete information (Binder et al., 2009; Fernandino et al., 2015; see Supplementary Figure 4). Thus, our results are in keeping with previous evidence showing two extended and separated sets of brain regions that support concrete and abstract knowledge (Wang et al., 2010).

However, contrary to some previous studies (Binder et al., 2009; Borghesani et al., 2016), we failed to find significant activity related to concrete features in the early visual cortex (EVC). One possible reason for this discrepancy is that, whereas these previous studies tracked the representation of single low-level visual features (e.g., size, color, movement), our concrete semantic predictor was derived from the dimensionality reduction of a multidimensional feature space spanning several sensory modalities (vision, touch, audition, etc.) and conceptual domains (color, face, music; see Supplementary Figure 5). This predictor is more likely to capture integrated sensorimotor information that is usually represented in convergence regions of the brain (Binder & Desai, 2011; Damasio, 1989). Indeed, the set of regions that in our analysis respond to the principal component of the concrete feature space (mPFC, Angular Gyrus, VOTC, IFG, SMG) largely coincide with regions that have been found to conjointly represent multiple concrete features (e.g., color, movement, motion, shape, manipulation) during conceptual retrieval (Fernandino et al., 2015). The anterior VOTC (see Figure 3E), which support the representation of several domains of knowledge (objects, faces, animals, etc.), in a format that is not tight to the visual modality only (Mattioni et al., 2020; Peelen & Downing, 2017; van den Hurk, Van Baelen, & Op de Beeck, 2017; Wang et al., 2015), qualifies as one of these convergence regions, in contrast with EVC that is highly modality-specific (Bottini et al., 2020; Wang et al., 2015).

Another interesting difference compared to some previous results is the involvement of the inferior frontal gyrus (IFG) in the representation of concrete features. Indeed, in some previous studies (including a meta-analysis), the IFG has been associated with the processing of abstract words, arguably for its role within the language system (Hoffman et al., 2015; Sabsevitz et al., 2005; Wang et al., 2010). Nevertheless, some studies before us reported IFG activation for concrete concepts, especially if related to actions (Kana, Blum, Ladden, & Ver Hoef, 2012; Rueschemeyer, Ekman, van Ackeren, & Kilner, 2014; Siri et al., 2008). It is important to consider that, in order to isolate brain regions dedicated to concrete and abstract feature representations, several previous studies contrasted different sets of concrete and abstract words. However, semantic reference is hard to control when creating different sets of words: Many abstract words may hold a strong reference to concrete features (e.g., sadness may be associated with darkness and freedom with a colorful and blooming landscape), and vice versa (a computer may be associated with intense cognitive and intellectual activity; Barsalou, Dutriaux, & Scheepers, 2018). One advantage of the current design, instead, is that the same words were contrasted based on their relative “location” in two multidimensional semantic spaces, one constituted by abstract and the other by concrete dimensions. Moreover, we controlled for several nuisance variables (e.g., word duration, frequency, response) by including them in the multiple regressions model and isolating brain activity that could be explained by these variables. These technical aspects may account for the differences between the current results and some previous studies that contrasted abstract with concrete words.

### Conclusions

We demonstrated that, during conceptual retrieval, abstract and concrete semantic information are represented in the ATL along a dorsal-to-medial gradient. During early stages of conceptual processing, right after lexical access, concrete and abstract features are encoded in posterior temporal regions and, at later time points, seem to converge into the ATL along separate cortical streams that coincide with long-range connectivity pathways leading to ATL subregions. Our timecourse analysis supports the hypothesis that the ventral-medial ATL receives concrete information from the ventral stream (related to object knowledge), and the dorsal-lateral ATL receives abstract information from the posterior STG and the orbito-frontal cortex (related to language processing and emotional value). In sum, we provided direct evidence that abstract and concrete semantic information travels along separate cortical routes during conceptual retrieval to reach the ATL gradient in later time windows, possibly coinciding with the retrieval of integrated and specific conceptual representations.

## Material and Methods

### Participants

Thirty native Italian speakers (11 female, aged 28.2 ± 4.8 years) participated in the study. All participants were right-handed and had no history of neurological or psychiatric disorders. Before testing participants gave their written informed consent and received monetary reimbursement for their participation. The experiment was conducted in accordance with the Declaration of Helsinki and was approved by the local ethical committee of the University of Trento.

### Experimental design

We derived our stimulus set from a previous work by Binder and colleagues (Binder et al., 2016). Out of 535 English words filed in Binder et al.’s (2016) original work, 438 were translated into Italian (352 nouns in the singular form, 54 verbs in the infinite tense and 32 adjectives in the singular male form). Selected words were 2 to 4 syllables long (M = 2.93, SD = 0.72) and could be unambiguously translated into Italian. Stimuli were recorded as 22050 Hz mono audio files, using the text-to-speech software ‘Speech2Go’ (SpeechWorks, Nuance communication, Burlington, MA, USA). Using Praat (Boersma & Weenink 2007), each audio file was trimmed of silence intervals at the beginning and at the end of the utterance and normalized to a uniform intensity. Finally, each file was inspected to detect acoustic anomalies or unnatural pronunciation.

Auditory stimuli were delivered via loudspeakers (Panphonics Sound Shower) placed inside the magnetically shielded MEG room. Stimuli were played at a comfortable sound level, which was the same for all participants. Stimulus presentation was controlled via Psychtoolbox (Brainard, 1997) running in a MATLAB 2015a environment.

Each trial started with 1.5s pre-stimulus silence followed by the auditory-presented word. Participants were instructed to categorize each stimulus as either related to sensory-perception (i.e. they express something that is related to one or more of the senses), or unrelated to sensory-perception. An auditory cue (a “beep” sound) prompted participants’ responses 2s after stimulus onset. Responses were collected via button presses operated with the dominant hand’s index and middle fingers. The response mapping was counterbalanced across participants. The maximum time given to respond was set to 2s and was followed by an interstimulus interval randomly jittered between 0.5 s and 1.0 s. Participants were familiarized with a short version of the task (30 trials taken from a different stimulus set) on a portable PC outside the MEG chamber. Participants were all blindfolded for the entire duration of the experiment and the room was kept in the dark. Each testing session lasted approximately 50 minutes and was divided into six, seven-minutes runs separated by short breaks.

### Uniqueness point

The time taken to access words’ lexical-semantic properties necessarily depends on the words themselves. The words used in the present study, for instance, varied greatly in length (M = 570.6ms; SD = 119.5ms); therefore aligning our analysis solely to the onset of each auditory stimulus would be suboptimal. We estimated the uniqueness point (UP) of each word. That is, the point in time when a string of sounds corresponds to one and only one word (Marlsen-Wilson, 1987). Ideally, this process would be carried out based on the phonological forms of words; however, phonological databases for Italian are very limited in size (e.g., PhonItalia, 120000 tokens; Goslin, Galluzzi, & Romani, 2014), which would have yielded imprecise estimates. Therefore, we took advantage of the near-perfect phoneme-to-grapheme correspondence in Italian, and computed UP based on orthographic databases, which are vastly larger (e.g., SUBTLEX–IT, 130M tokens; Crepaldi et al., 2013). The process can be summarized in two steps: 1) first, we divided the duration of each stimulus (auditory waveform) by the number of graphemes that constitute it. 2) The result was then multiplied by the orthographic UP (position in number of graphemes) of the lemmatized form of each stimulus (Goslin et al., 2013). The outcome of this procedure is the estimated position of the UP in each audio file. For the few words that clearly did not respect a clear phoneme-to-grapheme correspondence, this procedure was manually adjusted.

### MEG Data acquisition and preprocessing

MEG data were recorded using a whole-head 306 sensor (204 planar gradiometers; 102 magneto-meters) Vector-view system (Elekta Neuromag, Helsinki, Finland). Five head-position indicator coils (HPIs) were used to continuously determine the head position with respect to the MEG helmet. MEG signals were recorded at a sampling rate of 1 kHz and an online band-pass filter between 0.1 and 300 Hz. At the beginning of each experimental session, fiducial points of the head (the nasion and the left and right pre-auricular points) and a minimum of 300 other head-shape samples were digitized using a Polhemus FASTRAK 3D digitizer (Fastrak Polhemus, Inc., Colchester, VA, USA).

The raw data were processed using MaxFilter 2.0 (Elekta Neuromag ®). First, bad channels (identified via visual inspection) were replaced by interpolation. External sources of noise were separated from head-generated signals using a spatio-temporal variant of signal-space separation (SSS). Last, movement compensation was applied and each run was aligned to an average head position. All further analysis steps were performed in MATLAB 2019a using non-commercial software packages such as Fieldtrip (Oostenveld, Fries, Maris, & Schoffelen, 2011), Brainstorm (Tadel, Baillet, Mosher, Pantazis, & Leahy, 2011) and custom scripts. Continuous MEG recordings were filtered at 0.1 Hz using a two-pass Butterworth high-pass filter and epoched from -1.5 s before to 2s after the uniqueness point. Time segments contaminated by artifacts were manually rejected (total data lost of *M* = 7.6% SD = 7.7%). A Butterworth low-pass filter at 40Hz was applied to the epoched data. Before encoding, each trial segment was baseline corrected with respect to a 400ms time window before stimulus onset.

### Multiple linear regression analysis

Multiple linear regression analysis was applied to MEG data following the approach used in previous M/EEG studies (Chen et al., 2013, 2015; Hauk et al., 2006, 2009; Miozzo et al., 2015). The solution of a multiple regression provides the best least-square fit of all variables simultaneously to the data (Bertero, De Mol, & Pike, 1985). In M/EEG analysis the resulting event-related regression coefficients (ERRC) reflect the contribution of each predictor to the data for each time point, channel and subject. Importantly, because regression analysis is a special form of factorial designs, ERRC can be interpreted and analyzed as difference waves in ERP signals.

We focused on two semantic predictors, one abstract semantic component and one concrete semantic component (see below for details). All the regression models included psycholinguistic and word-form features as covariates (i.e., the word frequency, the duration of the word and the response given by the participant). Before encoding the predictors of each model were converted to normalized z-scores and tested for multicollinearity using a condition number test (Belsley et al., 1982). The output of the test is a condition index, which in the present study never exceeded a threshold of 4 (with values < 6 collinearity is not seen as a problem).

### Predictor variables

The aim of the present study was to investigate the contribution of abstract and concrete semantic dimensions of conceptual knowledge to single concepts representations. On this account, we derived our stimulus set from a previous work by Binder and colleagues (Binder et al., 2016). These authors collected ratings of the salience of 65 biologically plausible features to word meaning (for a detailed description of the procedure see Binder et al., 2016). For every word in the database (e.g., lemon) more than one thousand participants were asked to rate how associated was each of the features (e.g., color) with that aspect of the experience (e.g., would you define a lemon as having a characteristic or defining color?). The result is a semantic space where concepts can be represented as single entities into a multidimensional space having perceptual and conceptual features as dimensions. Crucially, features spanned both abstract and concrete domains of conceptual knowledge thus represent an ideal framework to operationalize our assumptions.

#### Semantic components

As mentioned above, more than sixty features composed our semantic space. Encoding of the entire space in one single model, however, would be suboptimal. In fact features are highly intercorrelated between each other, leaving us with a multicollinearity issue. One way this can be avoided is through dimensionality reduction techniques (Cunningham & Yu, 2014), such as principal component analysis (PCA). PCA generates a series of principal components (PCs) representing the same data in a new coordinate system, with the first PC usually accounting for the largest percentage of data variance. Following this rationale, we used PCA to derive two high-level semantic components. Three features (i.e., Complexity, Practice, Caused) were excluded due to incomplete ratings. An abstract component was obtained via PCA (after normalization) of 31 features encompassing Spatial, Temporal, Causal, Social, Emotion, Drive and Attention domains (the first PC explaining 27.4% of the overall variance). Similarly, the concrete component was calculated as the first principal component (24.7% of variance explained) of 31 features encompassing several sensory-motor domains such as Vision, Somatic, Audition Gustation, Olfaction and Motor domains.

#### Linguistic features

For each of the selected words, we obtained several psycholinguistic features: Word duration (M = 570ms, SD = 119ms) and Word Frequency (in Zipf’s scale, M = 4, SD = 0.8) were extracted from the SUBTLEX-IT database (Crepaldi et al., 2013); first syllable frequency (in the natural logarithm of token, M = 8.7, SD = 2.1) was extracted from PhonItalia (Goslin et al., 2014).

### Source reconstruction

Distributed minimum-norm source estimation (Hämäläinen & Ilmoniemi, 1994) was applied following the standard procedure in Brainstorm (Tadel et al., 2011). Anatomical T1-weighted MRI images were acquired during a separate session in a MAGNETOM Prisma 3T scanner (Siemens, Erlangen, Germany) using a 3D MPRAGE sequence, 1-mm^3^ resolution, TR = 2140ms, TI = 900ms, TE = 2.9ms, flip angle 12°. Anatomical MRI images were processed using an automated segmentation algorithm of the Freesurfer software (Fischl, 2012). Co-registration of MEG sensor configuration and the reconstructed scalp surfaces was based on ∼300 scalp surface locations. The data noise covariance matrix was calculated from the baseline interval of different predictors from the same model. The forward model was obtained using the overlapping spheres method (Huang, Mosher, & Leahy, 1999) as implemented in the Brainstorm software. Event-related regression coefficients were then projected onto a 15000 vertices boundary element using a dynamic statistical parametric mapping approach (dSPM; Dale et al., 2000). Dipole sources were assumed to be perpendicular to the cortical surface. Last, the individual results were projected to a default template (ICBM152) and spatially smoothed (3mm FWHM).

### ROIs

ROIs analysis was performed over three cortical areas and was restricted to the left hemisphere since the abstract-to-concrete gradient have never been found in the right ATL in previous distortion-corrected fMRI studies (Hoffman et al., 2015; Striem-Amit et al., 2018). Regions of interest included one early visual cortex (EVC) ROI (617 vertices), one ventral ROI (643 vertices) and one dorsal ROI (468 vertices) each combining brain areas adapted from the Desikan-Killiany cortical atlas. The EVC ROI consisted of the lateral occipital cortex, the calcarine fissure and the lingual gyrus. The ventral ROI encompassed regions of the ventral temporal-occipital cortex (i.e., the fusiform gyrus, the inferior temporal gyrus, the entorhinal cortex and the ventral-medial temporal pole). Conversely, the lateral orbitofrontal cortex (which was modified in order to exclude a small area protruding into the inferior frontal gyrus) and superior temporal gyrus composed the dorsal ROI. ROIs were designed following established models of white matter pathways connecting the dorsal-lateral and ventral-medial ATL with other cortical regions (Binney et al., 2012; Chen et al., 2017; and see Figure 3A). We divided the large ventral pathway (depicted in blue in Figure 3A) in EVC and VOTC because of their different role and level of processing during conceptual retrieval (Bottini et al., 2020; Bracci & Op de Beeck, 2016; Clarke & Tyler, 2014; Mattioni et al., 2020) and to obtain ROIs of comparable size. The orbito-frontal region and the STG were united in the same dorsal ROI because, according to the graded ATL model (Lambon-Ralph et al., 2017), both regions projects into the dorsal-lateral ATL and contribute to the representation of abstract features. Moreover, including OF and STG regions in a unique ROI allowed us to obtain regions of interest with a comparable number of vertices.

### Statistical analysis

In line with previous studies (Chen et al., 2013, 2015; Hauk et al., 2006; Miozzo et al., 2015) we depicted the time course of different regressors as the root-mean-square (RMS) of the signal-to-noise ratio (SNR) of ERRC. The SNR was computed on the grand mean of all subjects, by dividing the MEG signal at each channel and time point by the standard deviation of the baseline. This provided a unified (magnetometers and gradiometers are combined together) and easy-to-interpret measure of sensor-level activity. Source-reconstructed statistical maps of single predictors (e.g., Word frequency) were computed as consecutive 100ms average time-windows. Average signals within each time-window were tested against “0” using a whole-brain one-sample t-test (one tail), FDR corrected for multiple comparisons, *p* < .05 and a minimum number of vertices of 10. Source-reconstructed statistical maps of the contrast between predictors (e.g., Concrete > Abstract) were computed as consecutive 100ms average time-windows. Average signals within each time window were tested against “0” using a whole-brain one-sample t-test (two-tailed), FDR corrected for multiple comparisons, *p* < .05 and a minimum number of vertices of 10. Finally, ROI-constrained statistical maps for single predictors and contrast between predictors were computed as described above and restricting statistical comparisons within each ROI.

## Acknowledgments

We thank Virginie Crollen for helpful comments on an earlier version of this manuscript and Valentina Di Nunzio for help with the figures. The project was funded by a PRIN grant (Project number: 2015PCNJ5F_001) from the Italian Ministry of Education, University and Research (MIUR) awarded to Davide Crepaldi in collaboration with Olivier Collignon.

