## Supplementary materials for "Timecourse and convergence of abstract and concrete knowledge in the anterior temporal lobe"

#### Figure 1. Stimulus onset analysis

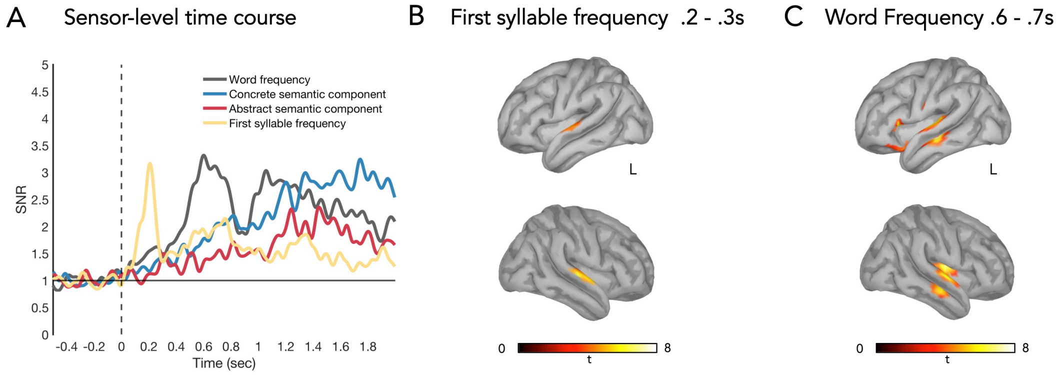


**Figure 1. Encoding of sublexical and lexical information. A) Root-mean-square of the SNR of ERRC of the first syllable frequency (yellow), word frequency (grey), concrete semantic component (blue) and abstract semantic component (red) predictors. 0s = stimulus onset. B) Source-reconstructed statistical maps of the First syllable frequency predictor (one-sample t-test (one tail), FDR-corrected *p* < .05, > 15-vertex) in the .2 to .3 interval. C) Source-reconstructed statistical maps of the Word Frequency predictor (one-sample t-test (one tail), FDR-corrected *p* < .05, > 15-vertex) in the .6 to .7 interval.**

#### Figure 2. Late word frequency effect

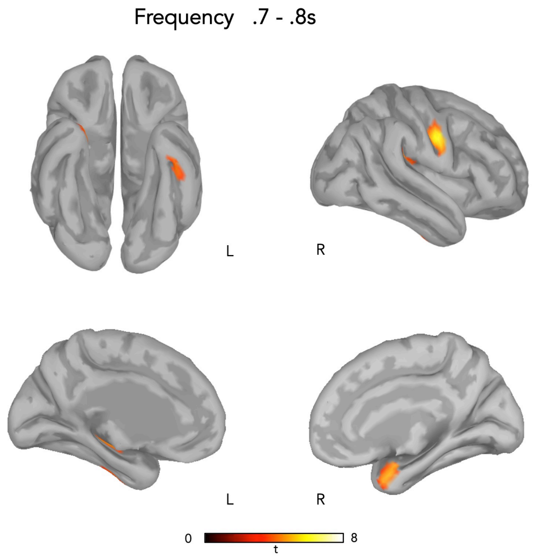


**Figure 2. Source-reconstructed statistical maps of the Word Frequency predictor (one-sample t-test (one tail), FDR-corrected *p* < .05, > 15-vertex) in the .7 to .8 interval.**

#### Figure 3. Superordinate gross categorical representations in the ATL

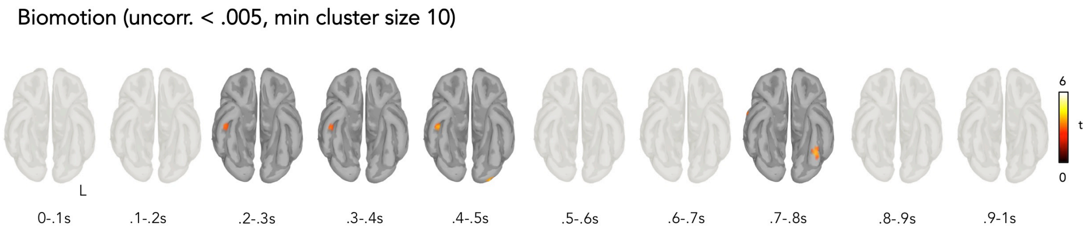


**Figure 3. Source-reconstructed statistical maps of the Biomotion predictor (one-sample t-test (one tail), uncorrected $p < .005$, > 10-vertex) in consecutive 100ms intervals.**

In order to test for superordinate gross categorical representations in the ATL we computed one additional model that included the feature Biomotion as well as nuisance variables (i.e., word duration, word frequency, participants' response). According to participants' ratings (Binder et al., 2016), the feature Biomotion is among the features that maximally distinguish between taxonomic categories such as Living Things vs. Artifacts (Figure 4 in Binder et al., 2016) and Animals vs. Tools (Figure 5 in Binder et al., 2016). We speculated that brain responses modulated by this feature reflect, at least to some degree, gross categorical representations (i.e., Living Things vs. Artifacts and Animals vs. Tools). Source-reconstructed statistical maps at early intervals (.2 - .5s) show that Biomotion was encoded, at a lower threshold (one-sample t-test (one tail), uncorrected $p < .005$, > 10-vertex), in the right ventral ATL. This suggests the possibility that these regions are related to semantic categorization processes as these ventral clusters correspond to the ones previously reported to be particularly sensitive to gross categorical distinctions (Borghesani, Buiatti, Eger, & Piazza, 2018; Chan et al., 2011; Teige et al., 2019). Given the explorative nature of this analysis, and the lack of significance with conventional threshold, this evidence should be considered with caution.

### Figure 4. Angular gyrus

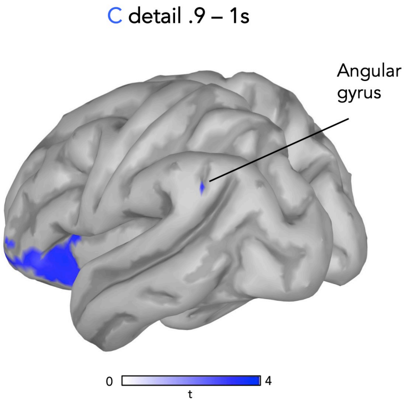


**Figure 4. Source-reconstructed statistical maps of the Concrete > Abstract predictor (one-sample t-test (one tail), FDR-corrected *p* < .05) in the .9 to 1s interval.**

### Figure 5. Concrete and Abstract features' space

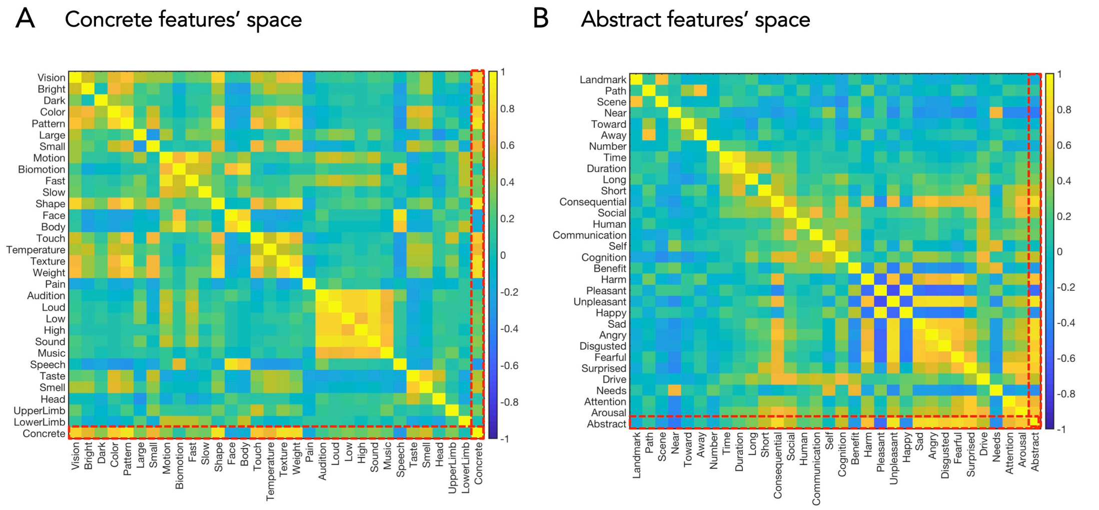


**Figure 5. Correlations of the concrete (A) and abstract (B) features' space. Red dotted boxes highlight concrete and abstract semantic components.**
